# New mutations, old statistical challenges

**DOI:** 10.1101/115964

**Authors:** Jeffrey C. Barrett, Joseph D. Buxbaum, David J. Cutler, Mark J. Daly, Bernie Devlin, Jacob Gratten, Matthew E. Hurles, Jack A. Kosmicki, Eric S. Lander, Daniel G. MacArthur, Benjamin M. Neale, Kathryn Roeder, Peter M. Visscher, Naomi R. Wray

## Abstract

Based on targeted sequencing of 208 genes in 11,730 neurodevelopmental disorder cases, Stessman et al. report the identification of 91 genes associated (at a False Discovery Rate [FDR] of 0.1) with autism spectrum disorders (ASD), intellectual disability (ID), and developmental delay (DD)—including what they characterize as 38 novel genes, not previously reported as connected with these diseases^1^.

If true, this would represent a substantial step forward. Unfortunately, each of the two discovery analyses (1. *De novo* mutation analysis and, 2. a comparison of private mutations with public control data) contain critical statistical flaws. When one accounts for these problems, fewer than half of the genes‐‐and very few, if any, of the novel findings‐‐survive. These errors have implications for how future analyses should be conducted, for understanding the genetic basis of these disorders, and for genomic medicine.

We discuss the two main analyses in turn and provide more detailed treatment of the issues in a supplementary technical note.

## 1. Two-stage analysis of *de novo* mutations

The authors selected 208 genes, consisting of 130 with one or more *de novo* truncating mutations in prior published studies, along with 78 others belonging to related pathways or having related Mendelian disease association. Of these genes, 19 have an already documented ‘genome-wide significant’ excess of *de novo* mutations from 5-6,000 neurodevelopmental disorder patients used as the “discovery sample” by the authors. Moreover, a more recently published exome dataset ^2^ has convincingly elevated the number of formally genome-wide significant genes to 93.

The authors resequenced this collection of 208 genes in 11,730 neurodevelopmental disorder cases (“replication sample”) and looked for an excess of *de novo* mutations. They report that analysis of *de novo* truncating mutations identifies 68 disease-associated genes, while analysis of *de novo* damaging missense mutations adds an additional 10 genes (with 32 of these 78 genes described as novel). These claims, however, are made based on a flawed statistical analysis, and we estimate fewer than half this number of genes achieve an FDR of 0.1, nearly all of which are known.

The study belongs to the traditional ‘two-stage’ design, which was first popularised 10-25 years ago, in early days of genome-wide association and, before that, linkage studies. There are two valid ways to analyze such a design: (1) count events only in the replication sample, in which case a significance threshold based on the number of genes tested in the replication sample (here, 208) is applied or (2) count events in both the initial and replication sample, in which case a significance threshold based on the number of genes in the genome (∼20,000) must be applied.

The authors, however, used an invalid approach: they counted events in both the initial and replication sample, but they applied the weaker significance threshold appropriate for testing only 208 genes. This approach inflates the significance of genes: because the 208 genes were chosen due to a higher-than-expected number of events in the initial sample, the number of events in the combined initial and replication sample will be artificially high; it will not follow the null distribution even if there are no associated genes. While seemingly subtle, this pernicious pitfall applies to any genomewide scanning technique: examining the first half of the data and then completing the study for only the most promising sites is still performing a full genome scan when both halves of the data are analyzed together.

The problem is underscored by some simple observations regarding the *de novo* likely gene disruptive (LGD) analysis:

1. The rate of *de novo* truncating mutations in the 208 genes is dramatically lower in the replication sample than in the initial sample (0.7% vs. 5.4%)
2. For 28 of the 68 genes claimed to be significant based on the presence of truncating *de novo* mutations, *no events at all* were observed in the replication sample. Despite the fact that the replication study (twice the size of the discovery sample) therefore provided evidence *against* these genes, the analysis declared the genes significant due only to the use of an inappropriately liberal significance level. These 28 genes include 11 of the ‘novel’ genes.

When one applies a correct significance threshold to the data for truncating *de novo* mutations, the number associated at FDR=0.1 falls from 68 genes to 30 genes - of which 23 reach the much stricter genome-wide significance threshold (Technical Note 1). As a general note we would encourage somewhat more emphasis on this stricter threshold than in many recent papers, with proper accounting for multiple variant categories tested, as the investment in clinical decision-making and functional studies should be informed by which genes have robust evidence and which may have a likely/probable but far from certain status.

Only *one* of the 30 genes, *SETBP1*, has not previously been reported in the exome literature (summarized in recent meta-analyses^2–6^) as associated with ASD, ID and/or DD, but this gene is a well-established cause of severe autosomal dominant intellectual disability, as published by some of the same authors in 2014 ^7^. Of the 29 others, 28 were documented in exome studies in 2015 or earlier and the 29th (*TCF4*) was noted in the most recent 2017 DDD paper but has been widely known as a cause of ID due to Pitt-Hopkins syndrome since 2007^8–10^.

## 2. Comparison of ‘private mutations’ in cases vs. public control data

The second type of analysis performed by the authors involves looking for an excess of “private mutations” (that is, singletons) in cases vs. controls. To assess significance, they use a permutation test in which they permute the labels of the cases and controls. Based on this analysis, the authors report 13 additional disease-associated genes (on top of the earlier 78).

This study design is statistically valid—*provided* that the cases and controls are chosen from the sample population and have been sequenced in the same manner. However, the results are *not* valid if the controls (i) did not come from the same populations (population history affects the entire allelic spectrum, including the frequency of singletons) and (ii) were not sequenced and analysed in identical ways (differences in the average sequence depth or the coverage of specific exons affects singleton detection). The use of a permutation test to assess significance does nothing to eliminate these problems: the systematic differences present in the case vs. control comparison are absent in the permutations which distribute the technical differences randomly.

In this case, however, the authors compared their case samples to a public control sample from the Exome Aggregation Consortium (ExAC)^11^ in a fashion that violated all three of the critical criteria above. The ExAC database is chosen from a different mix of populations and, being drawn from heterogeneous exome capture experiments, has differences in local and average coverage and is not at all matched to the targeted MIP study the authors use. For example, *UNC80* is highlighted as having an excess of private mutations in cases. However, examination of ExAC shows it is particularly poorly covered in most exome sequencing studies (in 58 of 64 *UNC80* exons fewer than half of ExAC samples achieve 20x depth)‐‐which will dramatically decrease the observed rate of singleton mutations.

To compare private mutations in cases vs. controls, it is essential that the samples be taken from the same populations, that the technical aspects of sequencing are matched across cases and controls, and that the authors are particularly careful to remove sites that are singleton in one study and are absent from the other study because the region was not sequenced or the variant was present at higher frequency and/or removed because of quality control filtering. Of note, the ExAC resource does provide summaries broken down by major continental ancestries and also provides coverage-depth information such that the data could be used much more accurately, albeit still imperfectly, in this context. Additionally, comparing the distributions of synonymous and other largely neutral variant categories should be useful for assessing the alignment of data sets sequenced and analyzed separately.

As presented in Stessman et al., the results from this analysis are not readily salvageable – even though many true positives may rise to the top of such an analysis, it is almost certain that false positives will also be present owing to the considerations outlined here. While the targeted nature of this study limits the number of such false positives, were the same strategy to be employed for comparison of whole-exome or genome data, the technical challenges would be quite considerable. In the companion technical note, both the theory and an empirical example are provided in support of these points. This analysis is conceptually sound and the field should be motivated to create shared genomic resources for which technical and population matching can be performed in a way to make these results reliable.

## Conclusion

Re-analysis suggests far fewer significant genes, and little or no truly novel genes relative to the existing literature, including the recent mega-analyses in ASD and DD/ID as well as well-established Mendelian genes. As the assignment of disease association to each gene is a foundational step on which years of molecular and clinical research will be built, it is absolutely imperative that studies of rare and *de novo* variation take heed of the hard-earned lessons and strict principles of the past decades of statistical genetics.

We urge the authors to update the analyses in this manuscript and generate corrected tables and figures reflecting an appropriate FDR correction in order to provide a more accurate view of per-gene association probabilities.

Signed (alphabetically)

---

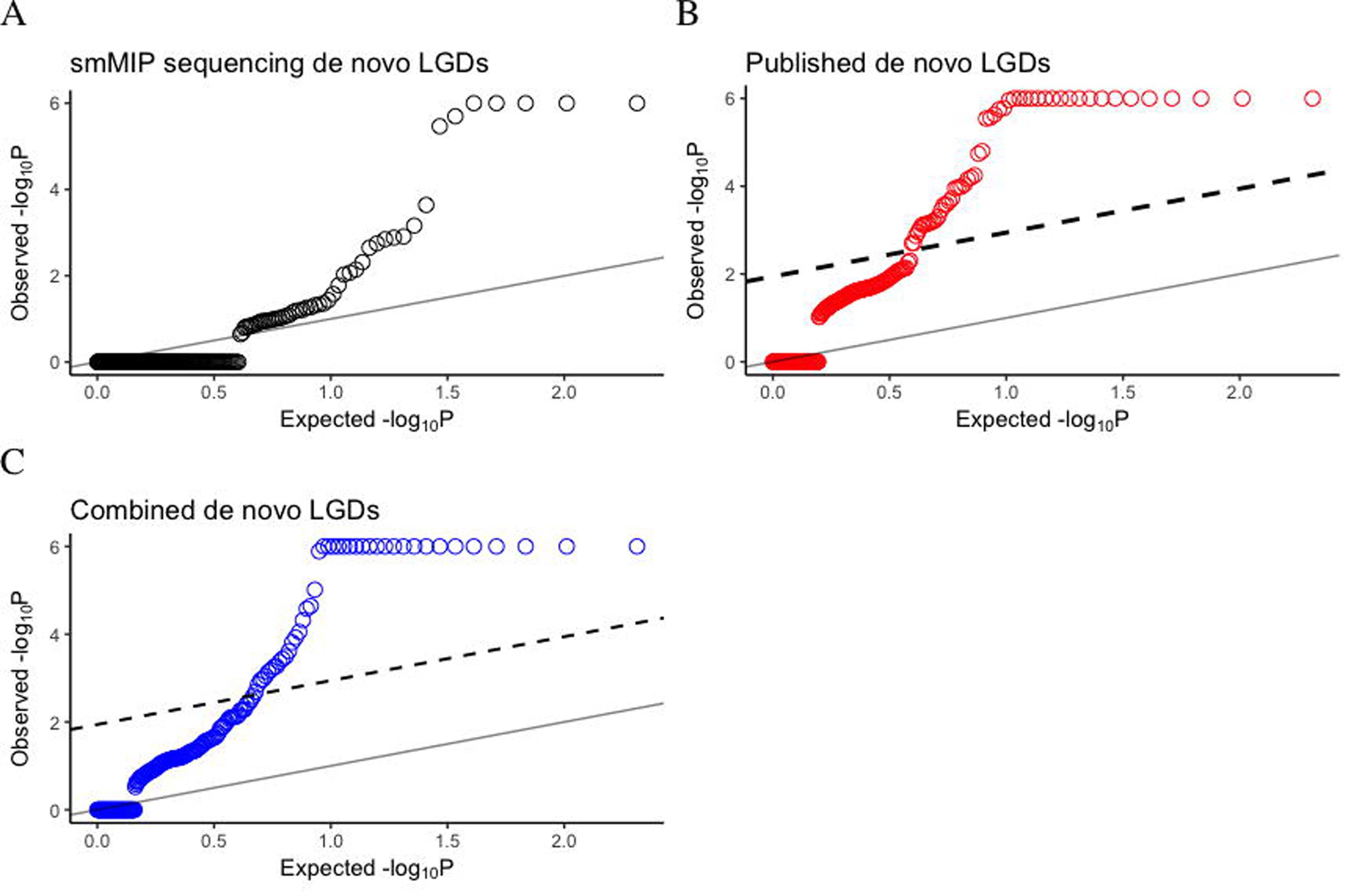

